# Assessing an age-old ecogeographical rule in nightjars across the full annual cycle

**DOI:** 10.1101/2023.08.30.555574

**Authors:** A Skinner, AM Korpach, S Åkesson, M Bakermans, TJ Benson, RM Brigham, GJ Conway, CM Davy, R Evens, KC Fraser, A Hedenström, IG Henderson, J Honkala, L Jacobsen, G Norevik, K Thorup, C Tonra, A Vitz, M Ward, E Knight

**Affiliations:** The University of British Columbia; University of Manitoba; Lund University; Worcester Polytechnic Institute; University of Illinois Urbana-Champaign; University of Regina; British Trust for Ornithology; Carleton University; University of Antwerp; Finnish Museum of Natural History; University of Copenhagen; The Ohio State University; Massachusetts Division of Fisheries and Wildlife; Alberta Biodiversity Monitoring Institute

## Abstract

Bergmann’s rule states that homeotherms are larger in colder climates (which occur at higher latitudes and elevations) due to thermoregulatory mechanisms. Despite being perhaps the most extensively studied biogeographical rule across all organisms, consistent mechanisms explaining which species or taxa adhere to Bergmann’s rule have been elusive. Furthermore, evidence for Bergmann’s rule in migratory animals has been mixed, and it was difficult to assess how environmental conditions across the full annual cycle impact body size until the recent miniaturization of tracking technology. Nightjars (Family Caprimulgidae), nocturnal birds with physiological and behavioral adaptations (e.g., torpor) to cope with the environmental extremes they often experience, offer a unique opportunity to elucidate the mechanisms underpinning Bergmann’s rule. Many nightjar species are strongly migratory and have large breeding ranges, offering the opportunity to look at variation in potential drivers within and across seasons of the annual cycle. Furthermore, variation in migration strategy within the family provides an opportunity to separate adaptations for migration strategy from adaptations for thermal tolerance. In this study, we use cross-continental data from three species of nightjars (Common nighthawk, Eastern whip-poor-will, and European nightjar) to assess 1) whether migratory species in this clade adheres to Bergmann’s rule, 2) which environmental factors are the best predictors of body size, and 3) the extent to which environmental conditions across the full annual cycle determine body size. For each species, we use breeding and winter location data from GPS tags to compare competing hypotheses explaining variation in body size: temperature regulation, productivity, and seasonality (during both the breeding and wintering periods), and migration distance. We found that Common nighthawk and Eastern whip-poor-will exhibit Bergmannian patterns in body size while European nightjar does not, although the spread of tag deployment sites on the breeding grounds was minimal for the European nightjar. Predictor variables associated with nightjar breeding locations more often explained body size than did variables on the wintering grounds. Surprisingly, models representing the geography hypothesis were best represented among important models in our final data set. Latitude and longitude correlated strongly with environmental variables and migratory distance; thus, these geographical variables offer a composite variable of sorts, summarizing many factors that likely influence body size in nightjars. Leveraging multi-species and cross-continental data across the full annual cycle, along with global environmental data, can provide insight into long-standing questions and will be important for understanding the generalizability of Bergmann’s rule.

## Introduction

Bergmann’s rule describes a pattern wherein ‘‘body size varies inversely with ambient temperature, so that body size increases with latitude,’’ (translated from the original German by Watt *et al*., 2010) The strong correlation between latitude and temperature implies that individuals of a species that tend to be larger in colder climates will indirectly be larger at higher latitudes (Salewski and Watt 2017). Some argue that Bergmann’s rule should explicitly include a thermoregulation mechanism (Watt *et al*., 2010a), whereas others suggest that observed patterns of latitudinal clines in body size can been called “Bergmannian” clines (Blackburn *et al*., 1999). Here, we use the term “Bergmann’s Rule” to refer to the pattern of latitudinal clines in body size, while recognizing that temperature variation is a likely mechanism. However, other environmental clines (e.g., productivity and seasonality) that covary with latitude can influence body size (Meiri, 2011). Therefore, relating patterns of body size variation to multiple potential environmental clines that covary with latitude can clarify the mechanism for the observed variation.

### Potential mechanisms for latitudinal clines in body size

Despite observed geographic clines in body size in most birds and mammals (Meiri & Dayan, 2003), there is still great uncertainty regarding the primary mechanisms. While an organism’s geographical position on the earth’s surface (latitude, longitude, and elevation) is often related to body size (i.e., the geography hypothesis), it is widely accepted that it is the underlying environmental clines that are highly correlated with earth’s geography and topography that exert selective pressures on body size. Numerous hypotheses have been examined to explain latitudinal patterns in body size (Blackburn *et al*., 1999; Jones *et al*., 2005). 1) The temperature regulation hypothesis was originally invoked by Bergmann (Watt *et al*., 2010b; Salewski & Watt, 2017), and suggested that homeothermic organisms closer to the poles experience selective pressure for efficient heat retention, and thus have larger body surface to body area ratios. This hypothesis has been expanded to also include more efficient heat dissipation in hotter climates (Blackburn *et al*., 1999). 2) The productivity hypothesis suggests that food availability is the most important determinant of body size, as organismal growth (particularly at the nestling stage) can be limited by nutrition (Rosenzweig, 1968; McNab, 2010; Yom-Tov & Geffen, 2011). Compelling evidence exists for this hypothesis, particularly where food availability and latitude are negatively related, and organismal body size follows gradients in food availability (larger at more southerly latitudes) (Katti & Price, 2003; Meiri *et al*., 2007). 3) The seasonality hypothesis (or the fasting endurance hypothesis; Blackburn *et al*., 1999; Ashton, 2002) suggests that larger animals are more resilient to periods of limited resources found in more seasonal environments (Lindstet & Boyce, 1985). This is because larger animals can store proportionally more fat and have relatively lower metabolic rates compared to their smaller-bodied conspecifics. High collinearity between the environmental variables representing these three distinct hypotheses has made it difficult to elucidate the mechanism behind latitudinal clines (Wigginton & Dobson, 1999; Yom-Tov & Geffen, 2011).

### Body size variation in migratory birds

Body size in both non-migratory and migratory bird species tend to follow latitudinal clines, but the relationship between latitude and body size is more complex in migratory species (Meiri & Dayan, 2003; Mainwaring & Street, 2021). Long-distance migratory bird species have body sizes and other morphological traits (wing shape and tail length) that are optimized for long flights (Hein *et al*., 2012; Phillips *et al*., 2018). Being light-weight facilitates energy-efficient long-distance flight, yet large wings, muscle hypertrophy, and fat stores also are important to provide the power and fuel for a costly journey (Alerstam *et al*., 2003). Wing size is also under strong selection from competing pressures including migration strategy (e.g., energy or time-efficient strategies), overall distance and route, foraging behavior (Marchetti 1995), and habitat associations across the full annual cycle (Rayner, 1988; Leisler & Winkler, 2003; Hedenström, 2008; Saino *et al*., 2017). Individuals of a species that migrate farther will likely have longer wings, but making predictions for body mass is challenging without a clear understanding of migratory strategy. While different combinations of behavioral and morphological adaptations exist to address the challenges of migration (Piersma, 2005; Dingle, 2006), it is important to consider the migration distance hypothesis (distinct from the “migration ability hypothesis” summarized in Blackburn *et al*., 1999), where body size is best determined by the constraints of migration (Gibson *et al*., 2019).

### Wintering grounds impacts of body size

Many species of migratory birds spend more than half of the year on the wintering grounds (La Sorte *et al*., 2017), and some studies have found environmental conditions on the wintering grounds to be important in determining body size as well (Gibson *et al*., 2019; Weeks *et al*., 2020). When this is the case, the body sizes observed across the breeding range (often where migratory species are measured) will be influenced by a species’ migratory connectivity, or the degree to which individuals that breed in close proximity to each other also overwinter in close proximity (Webster & Marra, 2005). Individuals from a species that exhibit weak migratory connectivity likely experience different winter conditions than other individuals from the same breeding population. Thus, morphology within breeding populations may be diverse if conditions on the overwintering grounds drive size variation in individuals. Strong migratory connectivity generally implies that individuals from a breeding population experience similar environmental conditions across the full annual cycle.

### Measures of body size

Most studies of latitudinal clines in body size use measures of wing length (Ashton, 2002). However, there is no single best metric of a bird’s body size, particularly for migratory birds. For example, body mass is ‘flexible’ through time (Gibson et al., 2019) because it is influenced by body condition or reproductive status, and can vary tremendously throughout the annual cycle, or even within a single day (Gosler *et al*., 1998). Wing length, on the other hand, is relatively ‘fixed’ in species that molt once a year (feather wear throughout the annual cycle negligibly influences wing length in most species) (e.g., Fernández & Lank, 2007). Wing length, therefore, is thought to be the best and most repeatable single measure of body size in birds (Gosler *et al*., 1998; Goodenough *et al*., 2010), but it can be confounded with selective pressures from migration, such as migration speed (Bennett *et al*., 2019), distance (Vágási *et al*., 2016), or strategy (Vincze *et al*., 2019). Given that the relationship between wing length and body mass is not isometric in all species (Rayner, 1988), using multiple measures of body size to test for Bergmannian clines may uncover previously undiscovered patterns or mechanisms.

### Nightjars as focal species

Nightjars (Family Caprimulgidae, hereafter ‘Caprimulgids’), with their unique life-history strategies, are an interesting focal group to understand how environmental pressures drive Bergmann’s rule in migratory species. Caprimulgids are largely crepuscular or nocturnal, and therefore are active at the coolest times of the day (Holyoak, 2001). However, many species roost or nest in open areas and experience high temperatures during the day on their breeding grounds (O’Connor *et al*., 2017; Newberry *et al*., 2021). Furthermore, Caprimulgids forage upon flying insects, whose activity is significantly reduced in cold weather (Taylor, 1963). Night temperatures may be quite cold at the northern extremes of breeding ranges, particularly when first arriving on their breeding grounds. Foraging activity is further restricted at certain times of the month when low moonlight levels inhibit aerial hunting by sight at night (Norevik *et al*., 2019; Evens *et al*., 2020; Souza-Cole, 2021), and thus, some individuals or species may have to endure long periods of fasting. Caprimulgids have therefore developed behavioral and physiological adaptations to withstand temperature extremes and periods of starvation, including low metabolic rates (Lane *et al*., 2004), torpor use (Brigham *et al*., 2006; Smit *et al*., 2011), gular fluttering, and efficient evaporative heat dissipation (O’Connor *et al*., 2017; Talbot *et al*., 2017; Newberry *et al*., 2021). Thus, Caprimulgids experience extreme environmental conditions, yet it is unclear the extent to which they rely on the thermodynamic properties of body size vs. behavioral and physiological adaptations to deal with these pressures.

We used temperate-breeding migratory Caprimulgids as a focal group to explore how environmental factors may drive latitudinal patterns in body size across the full annual cycle. We had the following objectives for this study: 1) determine whether migratory species from the Caprimulgid family adhere to the pattern of Bergmann’s rule (i.e., increasing size with latitude), 2) we test temperature, productivity, seasonality, and migration distance hypotheses to disentangle potential mechanisms of body size variation patterns in Caprimulgids, and 3) evaluate whether environmental conditions at the breeding or wintering grounds have a stronger influence on body size. We used cross-continental breeding and winter locations from three Caprimulgid species to fulfill the objectives: two Neotropical-Nearctic species (Eastern whip-poor-will – *Antrostomus vociferus* – and Common nighthawk – *Chordeiles minor*) and one palaearctic-Afrotropical species (European nightjar – *Caprimulgus europaeus*). Hereafter, these species will be referred to as the whip-poor-will, nighthawk, and European nightjar, respectively. We predicted (Table 1) that 1) birds breeding or wintering across larger latitudinal clines would adhere more strongly to Bergmann’s rule, because the range of environmental conditions they experience was likely greater; 2) migration distance would influence body size more strongly in long-distance migrants (nighthawk and European nightjar); and 3) environmental conditions on the breeding grounds would more strongly predict Bergmannian clines in body size than conditions on the wintering grounds, as Caprimulgids have to thermoregulate and provide resources to young as well (which may reduce their ability to rely upon behavioral adaptations).

**Table 1.** Nine potential hypotheses for the pattern / mechanism that explains Bergmann’s rule. We show the associated global model, and the predicted direction of effect for each variable: positive (+), negative (-), or differing predictions (+/-) by species or dependent variable. For example, for the migration distance hypothesis we predict that wing length will increase with longer migratory journeys, but make no prediction for body mass. Note the geography hypotheses examine patterns in body size, where all other hypotheses test the underlying mechanism. In addition to the introduction, see Blackburn et al. (1999) and Jones et al. (2005) for rationale behind hypotheses. CV = coefficient of variation.

| Hypothesis | Season | Global model |
| --- | --- | --- |
| Geography | Breeding | Latitude (+), Longitude (+/-), Elevation (+) |
| Geography | Winter | Latitude (+), Longitude (+/-), Elevation (+) |
| Temperature regulation | Breeding | Solar radiation (-), Temperature (-), Mean Daily Range (+/-) |
| Temperature regulation | Winter | Solar radiation (-), Temperature (-), Mean Daily Range (+/-) |
| Productivity | Breeding | Enhanced Vegetation Index (+), Precipitation (+) |
| Productivity | Winter | Enhanced Vegetation Index (+), Precipitation (+) |
| Seasonality | Breeding | Enhanced Vegetation Index CV (+), Precipitation CV (+), Temperature CV (+) |
| Seasonality | Winter | Enhanced Vegetation Index CV (+), Precipitation CV (+), Temperature CV (+) |
| Migration distance | No | Migratory distance (+/-) |

## Methods

### Capture, tag deployment, and measurement

All birds were captured using mist nets and conspecific playback in an initial deployment year (ranging from 2015 to 2021). We ringed birds using uniquely numbered aluminum bands, aged and sexed individuals, and measured wing chord using a wing ruler (unflattened in North America, flattened in Europe; precision = 1 mm) and mass using a digital scale (precision = 0.1 g), or a Pesola scale in rare cases (precision = 1.0 g). Whip-poor-wills and European nightjars were tagged with archival GPS loggers using a backpack style leg-loop harness (Rappole & Tipton, 1991), never exceeding 3.6% of the bird’s weight. Nighthawks were tagged with Argos-GPS loggers using a backpack harness (Åkesson et al. 2012) never exceeding 5.0% of the bird’s mass. Tags were scheduled to take a GPS fix (accuracy 10-m) every <1 – 10 days during migration and the over-winter period. We recaptured whip-poor-wills and European nightjars the following summer using the same methodology and expanded our search to neighboring territories (within 500-m) if the bird was not captured at the original site (recapture efforts from 2016 to 2022). Location data from the Argos-GPS tags on nighthawks was transmitted remotely via the Argos satellite system. See the following references for details relevant to deployment methods for each species (Knight *et al*., 2021 for nighthawks; Bakermans *et al*., 2022; Korpach *et al*., 2022; and Skinner *et al*., 2022b for whip-poor-wills; Lathouwers *et al*., 2022; and Norevik *et al*., 2019 for European nightjars).

### Body size estimation

We tested all hypotheses using either wing length or body mass as dependent variables. Generally, wing length increases proportionally with body mass to the one-third power (i.e., under isometric scaling), but this is species-specific and can vary substantially depending on a species’ life history. Whip-poor-wills and European nightjars exhibit near-isometric scaling between wing length and body mass, whereas scaling in nighthawks differs substantially from isometric (Fig S1). We considered using multivariate measures of body size (e.g., a principal component analysis), but the improvement from multivariate predictors is generally marginal (Gosler *et al*., 1998). Furthermore, this would have reduced sample sizes as each species and response variable combination had different sample sizes (whip-poor-will: mass n = 106, wing chord n = 100; nighthawk: mass n = 43, wing chord n = 35; European nightjar: mass n = 33, wing chord n = 29).

### Breeding & wintering location identification

To identify breeding and wintering ground locations, we used the location of capture for the breeding grounds and GPS data for the first wintering grounds (Fig 1). Some individuals had a second wintering site, but we used environmental information from the first wintering grounds location because 1) the majority of individuals had only one site, and 2) individuals that did move spent the majority of the overwinter period at their first non-breeding location (Norevik *et al*., 2017; Knight *et al*., 2021; Skinner, 2021; Bakermans *et al*., 2022; Korpach *et al*., 2022). If an individual was captured during multiple years, we discarded data from when individuals were juveniles, and averaged morphological data from all adult capture occasions. All individuals tracked across multiple years in our study wintered within a few hundred meters of previous years (16/16 of whip-poor-will, 7/7 European nightjars; Bakermans *et al*., 2022; Norevik *et al*., 2022a; Skinner *et al*., 2022b; Korpach et al., in prep), thus we used the winter location from the first year available. The majority of individuals caught across years on the breeding grounds were captured in the same location, and all individuals were captured within 2 km of the original capture location (Ng *et al*., 2018; Norevik *et al*., 2022a).

**Figure 1.**
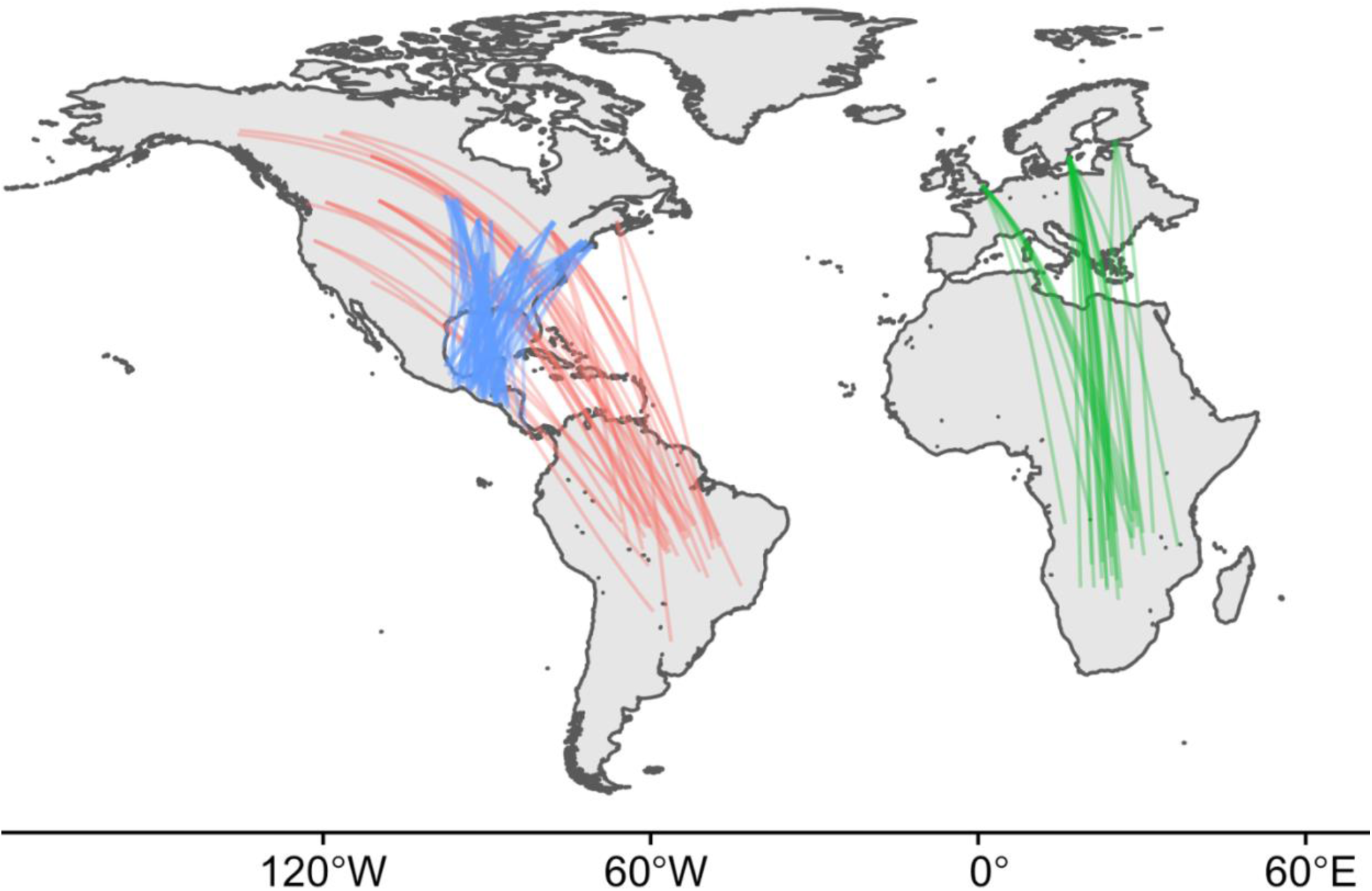
Breeding and wintering connections for 164 Caprimulgids tagged with archival GPS tags in North America and Europe (WGS 84). Lines do not represent actual migratory paths but instead the great-circle path between the breeding and wintering locations. Blue represents the Eastern whip-poor-will, red represents the Common nighthawk, and green represents the European nightjar.

### Environmental predictors

We calculated predictors using Google Earth Engine with the package *rgee* (Aybar, 2021) at both the breeding and wintering locations for each bird. We extracted climate variables from WorldClim 2.1 (historical climate averages from 1970-2000; Fick & Hijmans, 2017), and the Enhanced Vegetation Index (EVI) from Landsat 8 Collection Tier 1 8-Day EVI composites (Huete *et al*., 2002; Chander *et al*., 2009). We calculated the daily temperature range as the mean of each month’s max temperature - min temperature (sensu BIO2 from WorldClim’s bioclimatic variables). We calculated each variable as the mean value within a buffer that corresponded to the average home range size (rounded up to 1 significant figure) for each species in the corresponding season (whip-poor-will: breeding 500 m, winter 100 m; nighthawk: breeding 5000 m, winter 1000 m; European nightjar: 500 m, 100 m). See the following references for whip-poor-will (Wilson, 2003; Tonra *et al*., 2019; Bakermans *et al*., 2022; Skinner *et al*., 2022a), nighthawk (Ng *et al*., 2018), and European nightjar (Sharps *et al*., 2015; Mitchell *et al*., 2020). We used only the dates when individuals were thought to be stationary on the breeding or wintering grounds (breeding: May to September; winter: October to April). We averaged environmental variables or calculated the coefficient of variation (depending on the hypothesis in question) across the given months. We excluded highly correlated (|*r*| > 0.7; Fig S2) predictor variables from the same model and assessed variance inflation factors (VIF) with package *car* to ensure reasonable covariance of predictors in all models. Aside from one trivariate model in the nighthawk data set (for which VIF < 4.5 for both mass and wing), all VIF were ≤ 2.1.

### Statistical analysis

We organized the potential predictors of body size into unique and mutually exclusive hypotheses (Table 1). It is important to note that migration distance was calculated as the straight-line distance between breeding and wintering locations, and not the segmented migratory paths, which we did to avoid differences in temporal sampling resolution between projects. We modelled the relationship between body size (body mass or wing length) and our predictors using linear regression, restricting models to ≤ 3 variables each. We did not examine any interactions or polynomials due to small sample sizes. We examined the latitudinal distribution of ages (adult = 124, young = 12, unknown = 44) and sexes (male = 174, female = 12) across species, and ran linear models of body size against age or sex for each species. Body size only differed by age in whip-poor-wills (Wing chord: *F*_1,93_ = 5.09, *P* = 0.03; Mass: *F*_1,99_ = 4.94, *P* = 0.03), so we included age as a covariate in all whip-poor-will models. We scaled and centered all predictor and response variables to allow for model comparisons across species. We used a two-stage model selection process using Akaike’s information criterion corrected for small sample size (AIC_c_) to compare support for our hypotheses (Table 1).

In the first stage of model selection, we created 54 model sets (3 species × 2 response variables × 9 hypotheses). For each model set we created a global model and compared all possible subsets. Each model set had a null model to help determine the explanatory ability of the other models. The number of models (excluding the nulls) in each model set varied (between 2 and 8), as did the total number of models ran for each species (40 for Whip-poor-will, 49 for Common nighthawk, 45 for European nightjar). For each model set, we selected the most parsimonious model within ΔAIC_c_ ≤2 of the top model (Burnham & Anderson, 2004) for advancement to the next stage of model selection.

In the second stage of model selection, we compared the selected model from the first stage for each combination of species and response variable. This resulted in a single model set competing the 9 hypotheses (Table 1) for each species and response variable combination (6 final model sets). We considered the most parsimonious model within ΔAIC_c_ ≤2 in each set to be the “best” model, and models with ΔAIC_c_ ≤4 to be “important” models for addressing our first three objectives (Burnham & Anderson, 2004; Burnham *et al*., 2011). To address our first objective, we compared the best model of each set of candidate models to the null model using a likelihood ratio test to determine whether a species exhibited a Bergmannian pattern in body size and assess the goodness of fit of the model. To address the second and third objectives, we simply counted the number of times each hypothesis and season was represented in the final set of important models. We report results as mean ± standard error across all objectives.

## Results

### Predictors of Body Size

Nighthawks and whip-poor-wills generally exhibited patterns consistent with Bergmann’s rule (Fig 2) while European nightjars did not. In the final round of model selection there were 8 important models (ΔAIC_c_ ≤4) across both mass and wing chord, 4 for the nighthawk and 4 for the whip-poor-will (Table 2). Environmental pressures on the breeding grounds (n = 6 models) more often explained body size of Caprimulgids than pressures on the wintering grounds (n = 1) or migratory distance (n = 1). Geography models most often explained body size (n = 4 models), while models representing the other 4 hypotheses were present once each (Table 2). See Table S1 for results from the first round of model selection.

**Figure 2.**
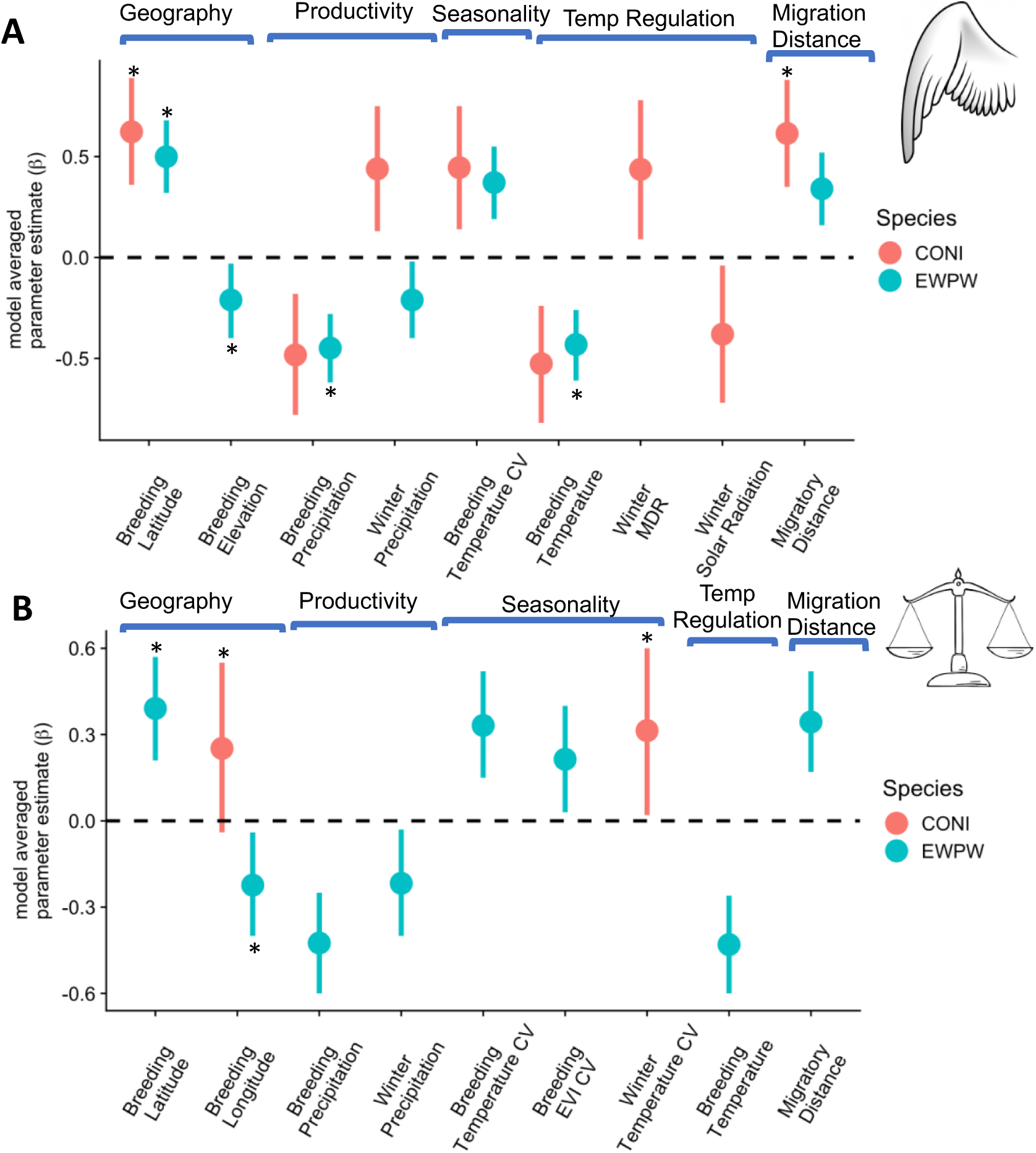
Parameter estimates and 95% unconditional confidence intervals of all variables that made it to the second round of model selection and their impact on body size for the Common nighthawk (CONI) and the Eastern whip-poor-will (EWPW). The European nightjar is not depicted as the null model was always within 2 ΔAIC_c_ points of the top models in the first round of model selection. Asterisks represent predictors of body size from the best overall models within ΔAICc ≤4 in round 2 of model selection. Both predictor and response variables were scaled and centered to facilitate direct comparison between species. Predictor variables are grouped by the underlying hypothesis they support (listed at the top of each panel). Panel a) depicts results where body size is approximated by wing chord, whereas in panel b) body size is approximated by body mass. CV is the coefficient of variation, MDR represents the mean daily temperature range, and EVI is the enhanced vegetation index.

**Table 2.**
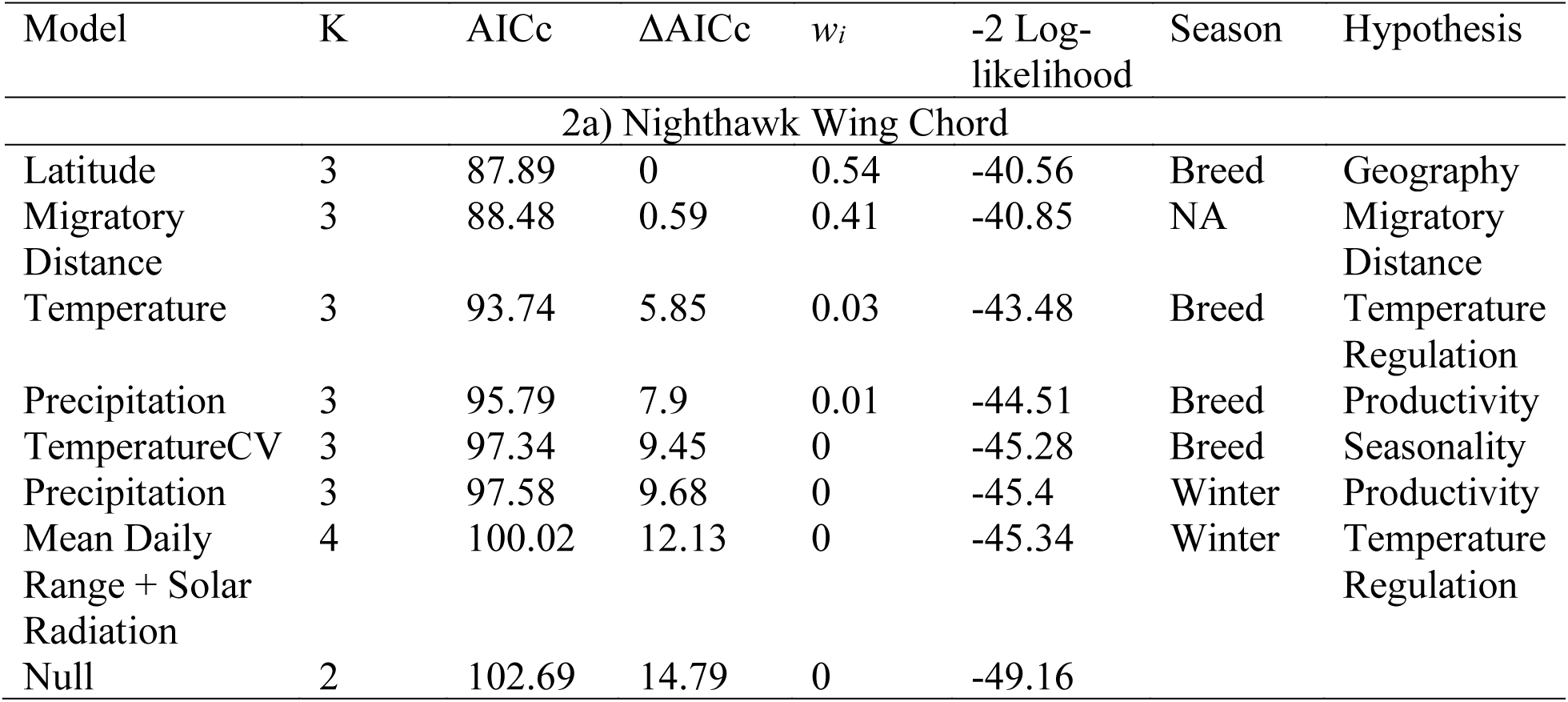
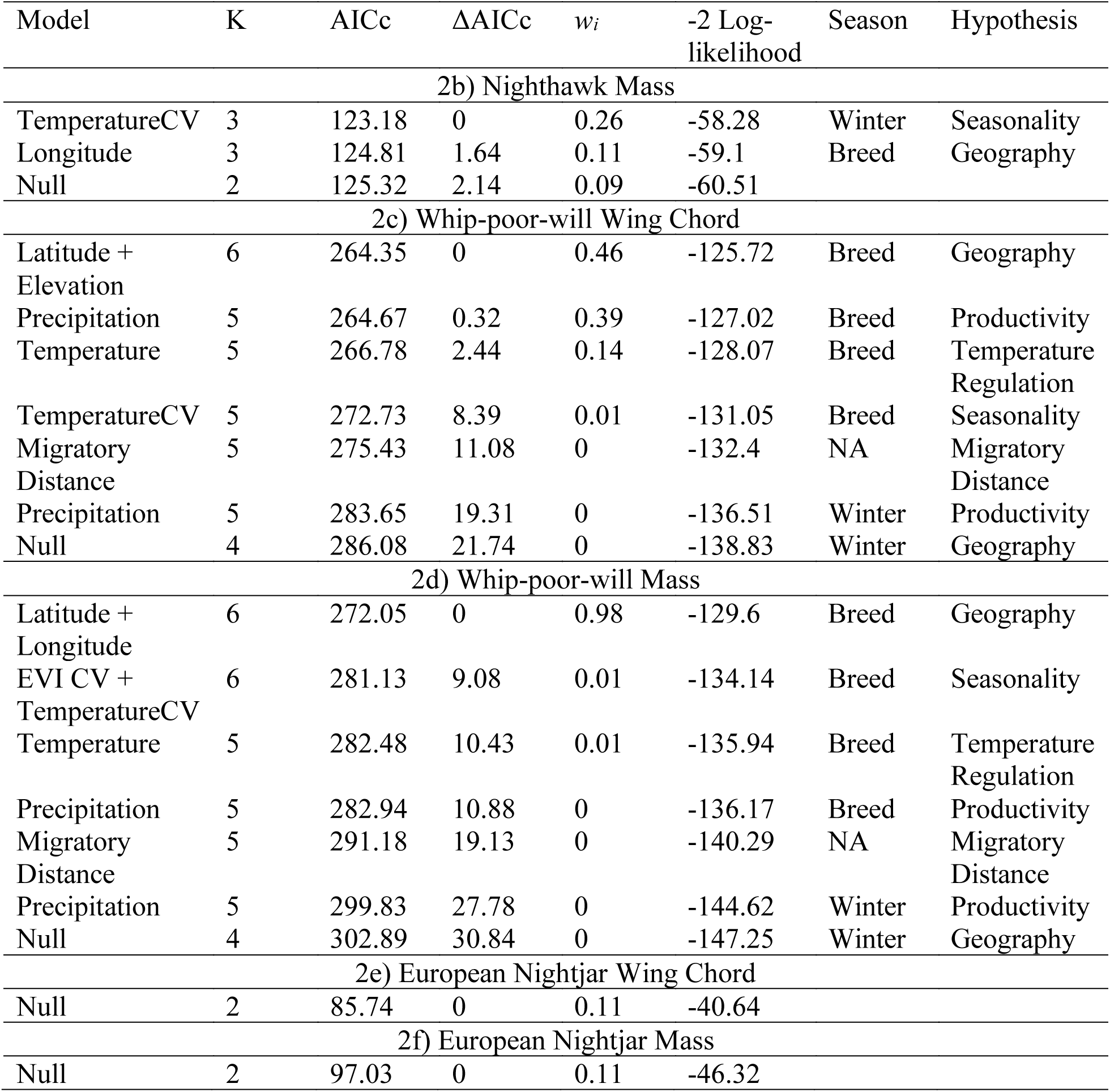
Final model sets for all species and response variable combinations. In Tables 2c and d (Whip-poor-will model sets), the categorical variable of age was included in all models. The total number of models in each set varied because the null model was often the most parsimonious model in round 1 of model selection. Models are ranked by the difference in Akaike’s information criterion corrected for small sample size (ΔAICc). Models with ΔAICc < 4 are considered “important”. K is the number of model parameters and *w_i_* is the model weight. Abbreviations are as follows: CV = coefficient of variation, and EVI = enhanced vegetation index.

The best model predicting nighthawk wing length included only breeding latitude (*w_i_* = 0.54), but the univariate model with migratory distance was also present amongst important models and received substantial support (*w_i_* = 0.41; Table 2). Wing length was longer as breeding latitude increased (β = 0.62, CI_95%_: 0.36 – 0.89; Fig 2). The best model predicting nighthawk mass included only the winter coefficient of variation (CV) of temperature, but breeding longitude was also present in important models (Table 2), although neither of these models received substantial Akaike’s weight (*w_i_* = 0.26 and *w_i_* = 0.11, respectively). Body mass increased with greater variation in winter temperature (β = 0.31, CI_95%_: 0.02 – 0.60). The best models for both dependent variables fit the data better than the null model (Wing: χ^2^ = 17.19, df = 1, *P* < 0.001; Mass: χ^2^ = 4.45, df = 1, *P* = 0.03).

The best model predicting whip-poor-will wing length included breeding latitude and breeding elevation (*w_i_* = 0.46), but models with breeding precipitation (*w_i_* = 0.39), as well as breeding temperature (*w_i_* = 0.14) were also present amongst important models. Whip-poor-will wing length was shorter with increased levels of breeding precipitation (β = -0.45 CI_95%_: -0.62 – -0.28). The only relevant model predicting whip-poor-will mass included breeding latitude and breeding longitude (*w_i_* = 0.98). Whip-poor-will mass increased in areas farther north (β = 0.39, CI_95%_: 0.21 – 0.57) and west (β = -0.22, CI_95%_: -0.40 – -0.04). The best models for both dependent variables fit the data better than the null model (Wing: χ^2^ = 23.63, df = 1, *P* < 0.001; Mass: χ^2^ = 35.29, df = 2, *P* < 0.001).

For the European nightjar, the null model was the best model across all hypotheses, although it is worth noting that the latitudinal spread of sampling locations across the breeding grounds was minimal (Fig 3).

**Figure 3.**
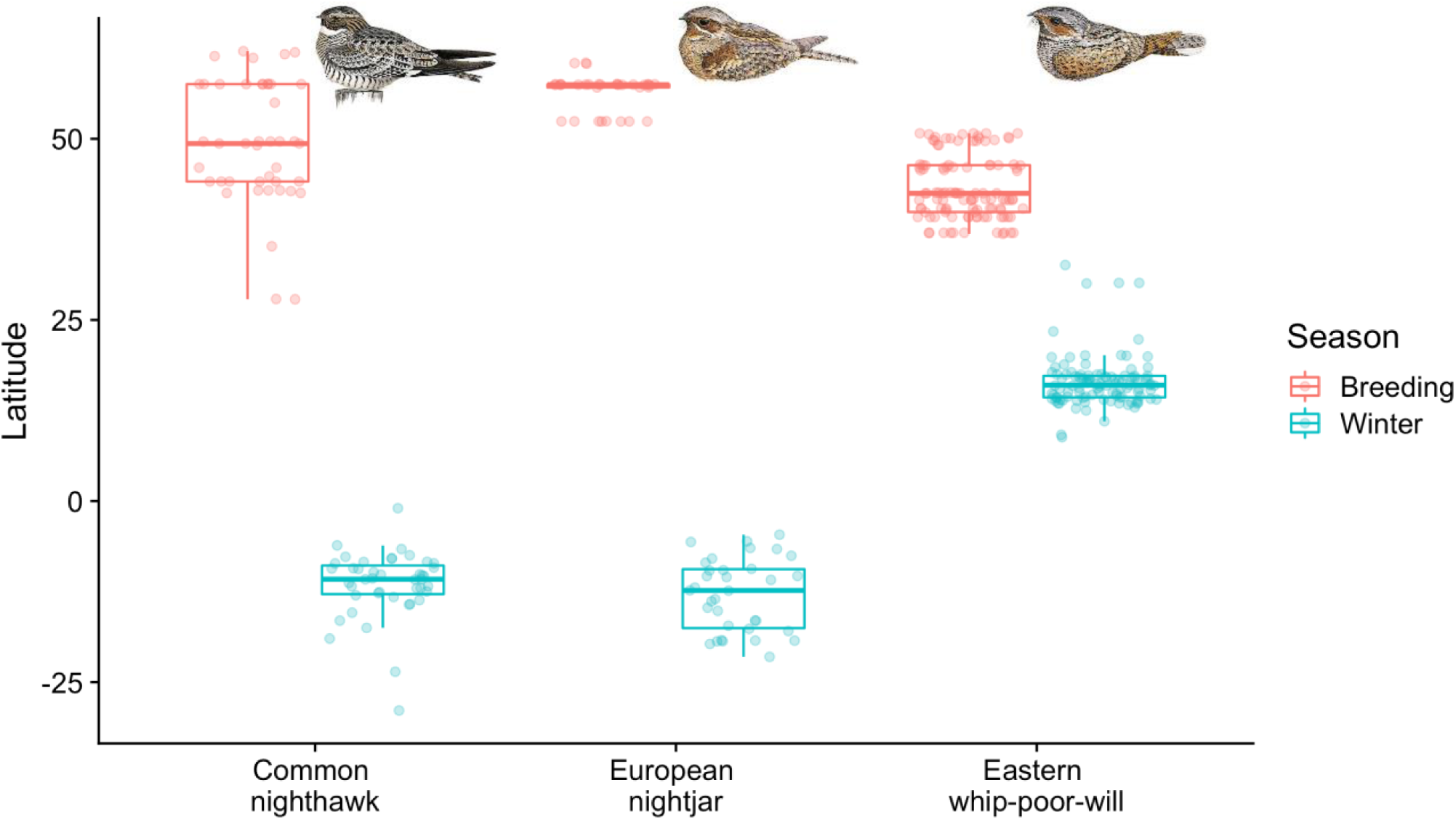
Researchers deployed tags across a greater range of latitudes in nighthawks (IQR = 13.4°; range = 34.2°) and whip-poor-wills (IQR = 6.5°; range = 13.9°) compared to the European nightjar (IQR = 0.4°; range = 8.1°). Most wintering locations (indicated by IQR) were more concentrated in space relative to the breeding grounds latitudes in nighthawks (winter: IQR = 4.0°; range = 27.9°) and whip-poor-wills (winter: IQR = 3.0°; range = 23.8°), but not European nightjar (winter: IQR = 8.1°; range = 16.9°). Absolute ranges were similar between breeding and wintering grounds, except for in the European nightjar which had greater variation in wintering grounds latitudes.

## Discussion

Bergmann’s rule is perhaps the oldest biogeographical rule Bergmann (1847), yet not all warm-blooded vertebrates follow this rule (Blackburn & Gaston, 1996; Ashton *et al*., 2000; Meiri & Dayan, 2003; Watt *et al*., 2010a), and a consensus regarding the underlying mechanism has evaded scientists. We leveraged cross-continental data from across the full annual cycle to understand whether Caprimulgids exhibit Bergmannian clines in body size and determine the most important factors driving those patterns. We also used the full annual cycle tracking data to subset only the relevant spatiotemporal scales for our independent variables, minimizing assumptions that were commonplace in the past. We found that whip-poor-wills and nighthawks both exhibited Bergmannian clines in body size, whereas European nightjars did not. In whip-poor-wills and nighthawks, variables associated with breeding locations were better predictors of body size than wintering ground locations, and geographic variables like latitude, longitude, and elevation were the best predictors of body size, although there was not resounding support for any single hypothesis. Despite being migratory, nocturnal, and having adaptations to mitigate extreme environmental conditions, Caprimulgids appear to exhibit similar patterns in body size variation as other groups of birds, adding support for the generalizability of Bergmann’s rule.

### Mechanisms underpinning Bergmann’s rule

Geographic variables were most frequently the best predictors of body size in nighthawks and whip-poor-wills (as in Gibson *et al*., 2019). Even Bergmann (1847), in his original rule, suggested that temperature underlies the latitudinal cline in body size. Thus, our findings are in contrast to previous studies that show body size tracking environmental gradients irrespective (or sometimes in the opposite predicted direction) of latitude (Katti & Price, 2003; Meiri *et al*., 2007; Nwaogu *et al*., 2018). In our study, latitude correlated strongly with environmental variables and migratory distance; thus, these geographical variables offer a composite variable of sorts, summarizing many factors that likely influence body size in Caprimulgids.

Surprisingly, however, breeding longitude and elevation were associated with Caprimulgid body size in the direction opposite to what we expected, at least in the whip-poor-will. Breeding longitude and precipitation were significantly positively correlated (i.e., precipitation increased farther east), and, amongst important models, breeding site precipitation was the only environmental variable that was opposite to the direction we predicted. Thus, the observed relationship between precipitation and size could also be responsible for the trend observed between longitude and size. We predicted that birds in rainier climates would be larger, because rainier climates tend to be richer in resources (i.e., the productivity hypothesis). However, we found that birds in more arid climates were larger. This finding is consistent with several previous studies, which implicated water regulation in the observed pattern (James, 1970; Nwaogu *et al*., 2018). Specifically, the relative amount of water lost to the environment is reduced in larger birds, thus favoring bigger individuals in arid climates. Thus, in combination with the fact that the Enhanced Vegetation Index did not appear in the second round of model selection, we find little support for the productivity hypothesis. It is noteworthy that breeding temperature also appeared in important models for whip-poor-will, adding to a growing body of evidence that both fluid and temperature regulation are critical in selecting for avian body size (James, 1970; Yom-Tov & Geffen, 2006; Nwaogu *et al*., 2018; Jirinec *et al*., 2021).

Whip-poor-will wing size also decreased as breeding elevation increased, which was opposite to what we predicted. Elevation was correlated with seasonality (e.g., CV of the Enhanced Vegetation Index), latitude, temperature, and longitude, yet elevation’s influence on body size was opposite to these other variables in all cases. For example, temperature decreased at higher elevations as expected, and overall whip-poor-wills had larger wings at cooler temperatures except at higher elevations where whip-poor-wills had smaller wings at higher elevations where it was cooler. Conversely, Youngflesh et al. (2022) found that wing length increased at higher elevations across (near-) passerines in North America and attributed this to the need to generate additional lift as air density is reduced at higher elevations. Thus, our findings with respect to elevation are surprising and contradict this recently proposed ecomorphological gradient. It is worth noting that elevation on the breeding grounds varied little (<500 m in whip-poor-wills) compared to other studies examining Bergmann’s rule across elevational gradients (e.g., Freeman, 2017; Youngflesh *et al*., 2022). Close examination of the trends opposite to what was predicted is critical as these can sometimes provide the best clues towards the true mechanisms underlying clines in body size.

### Environmental predictors across the full annual cycle

Variables at breeding locations were more important predictors of nighthawk and whip-poor-will body size than were variables at wintering sites. This may have occurred for three primary reasons. First, both experimental and observational evidence suggest that high temperatures during development lead to smaller nestlings (Andrew *et al*., 2018; Youtz *et al*., 2020). This implies that thermoregulation in the nestling period, before birds have grown feathers, may be particularly impactful on stress levels, survival, and ultimately in determining body size at the population level (Newberry & Swanson, 2018). This is likely exacerbated for in Caprimulgids, that nest in open habitats and are more exposed to environmental conditions relative to species that nest in closed habitats (Mainwaring & Street, 2021). Together, these studies and others (Speakman & Król, 2010; Andrew *et al*., 2017) imply that heat dissipation during hot breeding months may be more important than heat conservation in cold winter months. Second, it is possible that the responsibilities associated with chick-rearing (particularly feeding and thermoregulation) during the breeding period may prevent Caprimulgids from using evolved behavioral mechanisms for thermal tolerance (e.g., torpor), or strongly select for birds that can withstand periods of prolonged fasting (e.g., allowing for continued thermoregulation of chicks). Thus, Caprimulgids may rely more on behavioral approaches to maintain thermoregulatory and metabolic equilibrium in the non-breeding season. Third, all three Caprimulgids studied here had low migratory connectivity, such that birds from across large breeding ranges converge on core wintering areas (Norevik *et al*., 2020; Knight *et al*., 2021; Korpach *et al*., 2022; Skinner *et al*., 2022b). We expect winter environmental conditions to be fairly uniform in species with low migratory connectivity, although birds could select wintering locations at smaller spatial scales to provide more favorable environmental conditions for their given body size (especially if wintering in mountainous areas).

### Bergmann’s rule and climate change

An understanding of Bergmann’s rule is important for predicting species’ responses to climate change, and ultimately understanding which species are most vulnerable. As temperatures continue to warm globally, species’ long-term survival may depend upon the ability to shift their ranges northward or upslope, or evolve distinct morphologies better suited to deal with changing climates. There is evidence that nighthawk distribution may be shifting their breeding ranges northwards (see eBird [trend map](https://science.ebird.org/en/status-and-trends/species/comnig/trends-map)) (LaSorte & Thompson III, 2007), and climate change models predict that whip-poor-will ranges have the potential to expand at the north edge and contract at the south edge under a warming scenario (https://www.audubon.org/field-guide/bird/eastern-whip-poor-will). However, it is unclear whether their morphologies are also tracking environmental conditions through time, as seen by many bird species globally (Salewski *et al*., 2010; Weeks *et al*., 2020; Jirinec *et al*., 2021). Generally, bird morphology evolves quickly (Egbert & Belthoff, 2003), and additive models are often required to adequately model the rapid change in morphological response to temperature change through time (e.g., Weeks *et al*., 2020). However, the mismatch we observed between morphology and environmental conditions experienced on the wintering grounds is concerning and may be part of the reason these three species are declining. Climate change may further exacerbate this mismatch, particularly given that Caprimulgids appear to have high site fidelity across the annual cycle (Ng *et al*., 2018; Bakermans *et al*., 2022; Norevik *et al*., 2022b) and long lifespans (> 10 year), resulting in a lag between range shifts and climate shifts. Climate change appears to most strongly impact long-distance migrants that exhibit low diversity in migratory strategy (e.g., obligate migrants, small non-breeding ranges; Both *et al*., 2010; Gilroy *et al*., 2016), as is the case of the three Caprimulgids examined in this study. An understanding of how Caprimulgids respond to a changing climate throughout the full annual cycle will be critical to their conservation.

### Future directions for assessing Bergmann’s rule

Old-world species, including other groups of nocturnal birds, generally adhere to Bergmann’s rule (Ashton, 2002, Romano *et al*., 2021). Therefore, given the similarities in life-history between European nightjars and the two new-world species we examined, we surmise that we did not detect a Bergmannian cline in European nightjars because of the limited sampling extent. Most European nightjars (70%) in our study were tagged at two sites in Sweden that are <0.5° latitude apart, thus greatly minimizing our ability to detect a Bergmannian cline in body size on the breeding grounds. Given that European nightjars wintered across a relatively large portion of Africa (IQR of winter latitudes was >8°), it is unlikely that environmental conditions on the wintering grounds exhibit a strong influence on body size. However, an important next step is to leverage a larger data set of existing European nightjar data across the breeding grounds to increase the ability to detect an effect in this phase of the annual cycle.

Focal systems with low collinearity among environmental covariates and high migratory connectivity are ideal for assessing the drivers of body size in species; however, these ideal qualities are difficult to find. The rare cases when environmental predictors vary with latitude in the direction opposite to what is expected are particularly valuable (e.g., Katti & Price, 2003; Meiri *et al*., 2007), as are manipulative studies (e.g., Andrew *et al*., 2017) and meta-analyses (e.g., Ashton, 2002). Our study also illustrates the importance of using multiple metrics of body size, particularly in migratory species. Correlations between wing size and migratory distance are likely to be strong in most species, and mass can fluctuate throughout the annual cycle due to changes in muscle and fat composition (Liknes & Swanson, 2011). Thus, there are issues with either metric when used in isolation. Furthermore, it is important to understand the relationship between linear measurements and mass; allometric scaling theory provides the exploratory framework for understanding these relationships and must be utilized for a nuanced understanding of results.

### Conclusions

We found that two of the three Caprimulgids we studied adhered to Bergmann’s rule, and that explanatory variables associated with the breeding grounds, particularly related to the bird’s geographical position in space, were the best predictors of body size. We did not find conclusive evidence for any one environmental hypothesis and suggest that geographic variables likely rose to the top in model selection because they combine information across many important environmental gradients. In migratory organisms, Bergmann’s rule has often been assessed with the most easily accessible data – geographic information from the breeding grounds (Ashton, 2002). Our research suggests that these readily available data may indeed represent the most important variables influencing body size, and thus, past conclusions on Bergmann’s rule in migratory birds may be robust, although this assumption should be examined in a greater suite of species. Ultimately, the current era of remote sensing, readily available environmental data, and increasingly affordable tracking technology make it an exciting time to better understand Bergmann’s rule in migratory species.

