## Supplemental information for "Assessing an age-old ecogeographical rule in nightjars across the full annual cycle"

Table S1. Initial (round 1) model sets for all species and response variable combinations, where each model set represents a hypothesis for a given season (e.g., geography in the breeding grounds; n = 54 model sets). In Whip-poor-will model sets, the categorical variable of age is not listed but was included in all models. Models are ranked by the difference in Akaike's information criterion corrected for small sample size (ΔAICc). K is the number of model parameters and $w_i$ is the model weight. Abbreviations are as follows: CV = coefficient of variation, and EVI = enhanced vegetation

| Modnames | K | AICc | ΔAICc | $w_i$ | -2 Log-likelihood |
|---|---|---|---|---|---|
| CONI-Wing.Chord-Breed-Geo | | | | | |
| Latitude | 3 | 87.89 | 0 | 0.52 | -40.56 |
| Latitude + Longitude | 4 | 89.32 | 1.42 | 0.25 | -39.99 |
| Latitude + Elevation | 4 | 90.45 | 2.56 | 0.14 | -40.56 |
| Latitude + Longitude + Elevation | 5 | 91.77 | 3.88 | 0.07 | -39.85 |
| Longitude | 3 | 98.02 | 10.13 | 0 | -45.62 |
| Longitude + Elevation | 4 | 98.65 | 10.76 | 0 | -44.66 |
| 1 | 2 | 102.69 | 14.79 | 0 | -49.16 |
| Elevation | 3 | 105.04 | 17.15 | 0 | -49.14 |
| CONI-Wing.Chord-Winter-Geo | | | | | |
| 1 | 2 | 102.69 | 0 | 0.3 | -49.16 |
| Elevation | 3 | 103.5 | 0.82 | 0.2 | -48.36 |
| Longitude | 3 | 104.25 | 1.57 | 0.14 | -48.74 |
| Latitude | 3 | 104.66 | 1.97 | 0.11 | -48.94 |
| Latitude + Elevation | 4 | 104.94 | 2.25 | 0.1 | -47.8 |
| Latitude + Longitude | 4 | 105.72 | 3.04 | 0.07 | -48.2 |
| Longitude + Elevation | 4 | 106.06 | 3.38 | 0.06 | -48.36 |
| Latitude + Longitude + Elevation | 5 | 107.6 | 4.91 | 0.03 | -47.76 |
| CONI-Wing.Chord-Breed-TR | | | | | |
| Temperature | 3 | 93.74 | 0 | 0.39 | -43.48 |

| | | | | | |
|---|---|---|---|---|---|
| Solar Radiation + Temperature | 4 | 95.2 | 1.46 | 0.19 | -42.93 |
| Mean Daily Range + Solar Radiation | 4 | 95.25 | 1.51 | 0.19 | -42.96 |
| Mean Daily Range + Temperature | 4 | 96.19 | 2.45 | 0.12 | -43.43 |
| Mean Daily Range + Solar Radiation + Temperature | 5 | 97.01 | 3.27 | 0.08 | -42.47 |
| Solar Radiation | 3 | 98.83 | 5.08 | 0.03 | -46.03 |
| 1 | 2 | 102.69 | 8.95 | 0 | -49.16 |
| Mean Daily Range | 3 | 104.82 | 11.08 | 0 | -49.02 |
| CONI-Wing.Chord-Winter-TR | | | | | |
| Mean Daily Range + Solar Radiation | 4 | 100.02 | 0 | 0.44 | -45.34 |
| Mean Daily Range | 3 | 102.29 | 2.27 | 0.14 | -47.76 |
| 1 | 2 | 102.69 | 2.67 | 0.12 | -49.16 |
| Mean Daily Range + Solar Radiation + Temperature | 5 | 102.75 | 2.73 | 0.11 | -45.34 |
| Solar Radiation | 3 | 103.71 | 3.69 | 0.07 | -48.47 |
| Mean Daily Range + Temperature | 4 | 104.69 | 4.67 | 0.04 | -47.68 |
| Temperature | 3 | 104.9 | 4.88 | 0.04 | -49.06 |
| Solar Radiation + Temperature | 4 | 105.34 | 5.33 | 0.03 | -48.01 |
| CONI-Wing.Chord-Breed-Prod | | | | | |
| Precipitation | 3 | 95.79 | 0 | 0.73 | -44.51 |
| EVI + Precipitation | 4 | 98.26 | 2.48 | 0.21 | -44.47 |
| EVI | 3 | 101.61 | 5.82 | 0.04 | -47.42 |
| 1 | 2 | 102.69 | 6.9 | 0.02 | -49.16 |

| | | | | | |
|---|---|---|---|---|---|
| CONI-Wing.Chord-Winter-Prod | | | | | |
| Precipitation | 3 | 97.58 | 0 | 0.71 | -45.4 |
| EVI + Precipitation | 4 | 100.02 | 2.45 | 0.21 | -45.34 |
| 1 | 2 | 102.69 | 5.11 | 0.06 | -49.16 |
| EVI | 3 | 104.19 | 6.61 | 0.03 | -48.71 |
| CONI-Wing.Chord-Breed-Seas | | | | | |
| TemperatureCV | 3 | 97.34 | 0 | 0.57 | -45.28 |
| EVI CV + TemperatureCV | 4 | 98.23 | 0.89 | 0.37 | -44.45 |
| 1 | 2 | 102.69 | 5.35 | 0.04 | -49.16 |
| EVI CV | 3 | 104.21 | 6.87 | 0.02 | -48.72 |
| CONI-Wing.Chord-Winter-Seas | | | | | |
| 1 | 2 | 102.69 | 0 | 0.45 | -49.16 |
| EVI CV | 3 | 103.66 | 0.97 | 0.28 | -48.44 |
| TemperatureCV | 3 | 104.34 | 1.65 | 0.2 | -48.78 |
| EVI CV + TemperatureCV | 4 | 106.1 | 3.41 | 0.08 | -48.38 |
| CONI-Wing.Chord-NA-Mig.Dist | | | | | |
| Migratory Distance | 3 | 88.48 | 0 | 1 | -40.85 |
| 1 | 2 | 102.69 | 14.21 | 0 | -49.16 |
| CONI-Mass-Breed-Geo | | | | | |
| Latitude + Longitude + Elevation | 5 | 123.1 | 0 | 0.33 | -55.74 |
| Latitude + Longitude | 4 | 123.55 | 0.45 | 0.26 | -57.25 |
| Longitude | 3 | 124.81 | 1.71 | 0.14 | -59.1 |
| 1 | 2 | 125.32 | 2.22 | 0.11 | -60.51 |
| Longitude + Elevation | 4 | 126.46 | 3.36 | 0.06 | -58.7 |
| Latitude | 3 | 127.12 | 4.01 | 0.04 | -60.25 |
| Elevation | 3 | 127.58 | 4.48 | 0.04 | -60.48 |
| Latitude + Elevation | 4 | 129.46 | 6.35 | 0.01 | -60.2 |

| | | | | | |
|---|---|---|---|---|---|
| CONI-Mass-Winter-Geo | | | | | |
| 1 | 2 | 125.32 | 0 | 0.22 | -60.51 |
| Latitude | 3 | 125.59 | 0.28 | 0.2 | -59.49 |
| Longitude + Elevation | 4 | 126.34 | 1.02 | 0.13 | -58.64 |
| Longitude | 3 | 126.52 | 1.2 | 0.12 | -59.95 |
| Latitude + Longitude + Elevation | 5 | 127.06 | 1.74 | 0.09 | -57.72 |
| Latitude + Elevation | 4 | 127.35 | 2.04 | 0.08 | -59.15 |
| Elevation | 3 | 127.48 | 2.17 | 0.08 | -60.43 |
| Latitude + Longitude | 4 | 127.58 | 2.27 | 0.07 | -59.27 |
| CONI-Mass-Breed-TR | | | | | |
| 1 | 2 | 125.32 | 0 | 0.28 | -60.51 |
| Temperature | 3 | 125.41 | 0.1 | 0.27 | -59.4 |
| Solar Radiation | 3 | 127.01 | 1.69 | 0.12 | -60.2 |
| Mean Daily Range | 3 | 127.61 | 2.3 | 0.09 | -60.5 |
| Mean Daily Range + Temperature | 4 | 127.82 | 2.5 | 0.08 | -59.38 |
| Solar Radiation + Temperature | 4 | 127.85 | 2.53 | 0.08 | -59.4 |
| Mean Daily Range + Solar Radiation | 4 | 128.98 | 3.67 | 0.05 | -59.97 |
| Mean Daily Range + Solar Radiation + Temperature | 5 | 130.27 | 4.95 | 0.02 | -59.32 |
| CONI-Mass-Winter-TR | | | | | |
| Solar Radiation | 3 | 124.19 | 0 | 0.36 | -58.79 |
| 1 | 2 | 125.32 | 1.12 | 0.21 | -60.51 |
| Mean Daily Range + Solar Radiation | 4 | 126.63 | 2.43 | 0.11 | -58.79 |
| Solar Radiation + Temperature | 4 | 126.63 | 2.43 | 0.11 | -58.79 |
| Mean Daily Range | 3 | 127.09 | 2.9 | 0.08 | -60.24 |
| Temperature | 3 | 127.25 | 3.06 | 0.08 | -60.32 |

| | | | | | |
|---|---|---|---|---|---|
| Mean Daily Range + Solar Radiation + Temperature | 5 | 129.2 | 5 | 0.03 | -58.79 |
| Mean Daily Range + Temperature | 4 | 129.43 | 5.24 | 0.03 | -60.19 |
| CONI-Mass-Breed-Prod | | | | | |
| 1 | 2 | 125.32 | 0 | 0.36 | -60.51 |
| Precipitation | 3 | 125.83 | 0.52 | 0.28 | -59.61 |
| EVI | 3 | 125.93 | 0.61 | 0.27 | -59.66 |
| EVI + Precipitation | 4 | 127.98 | 2.67 | 0.1 | -59.47 |
| CONI-Mass-Winter-Prod | | | | | |
| 1 | 2 | 125.32 | 0 | 0.42 | -60.51 |
| EVI | 3 | 126.1 | 0.78 | 0.28 | -59.74 |
| EVI + Precipitation | 4 | 127.22 | 1.9 | 0.16 | -59.08 |
| Precipitation | 3 | 127.49 | 2.17 | 0.14 | -60.44 |
| CONI-Mass-Breed-Seas | | | | | |
| TemperatureCV | 3 | 125.08 | 0 | 0.4 | -59.23 |
| 1 | 2 | 125.32 | 0.23 | 0.35 | -60.51 |
| EVI CV + TemperatureCV | 4 | 127.39 | 2.31 | 0.13 | -59.17 |
| EVI CV | 3 | 127.41 | 2.33 | 0.12 | -60.4 |
| CONI-Mass-Winter-Seas | | | | | |
| TemperatureCV | 3 | 123.18 | 0 | 0.4 | -58.28 |
| EVI CV | 3 | 124.07 | 0.89 | 0.26 | -58.73 |
| EVI CV + TemperatureCV | 4 | 124.48 | 1.3 | 0.21 | -57.71 |
| 1 | 2 | 125.32 | 2.14 | 0.14 | -60.51 |
| CONI-Mass-NA-Mig.Dist | | | | | |
| 1 | 2 | 125.32 | 0 | 0.76 | -60.51 |
| Migratory Distance | 3 | 127.59 | 2.28 | 0.24 | -60.49 |
| EWPW-Wing.Chord-Breed-Geo | | | | | |
| Latitude + Elevation | 6 | 264.35 | 0 | 0.5 | -125.72 |

| | | | | | |
|---|---|---|---|---|---|
| Latitude + Longitude | 6 | 264.9 | 0.55 | 0.38 | -126 |
| Latitude | 5 | 267.18 | 2.83 | 0.12 | -128.27 |
| 1 | 4 | 286.08 | 21.74 | 0 | -138.83 |
| Elevation | 5 | 288.16 | 23.81 | 0 | -138.76 |
| Longitude | 5 | 288.2 | 23.85 | 0 | -138.78 |
| Longitude + Elevation | 6 | 290.41 | 26.07 | 0 | -138.76 |
| EWPW-Wing.Chord-Winter-Geo | | | | | |
| 1 | 4 | 286.08 | 0 | 0.4 | -138.83 |
| Elevation | 5 | 287.71 | 1.63 | 0.18 | -138.54 |
| Latitude | 5 | 287.8 | 1.71 | 0.17 | -138.58 |
| Longitude | 5 | 288.24 | 2.16 | 0.13 | -138.8 |
| Latitude + Elevation | 6 | 289.79 | 3.7 | 0.06 | -138.44 |
| Longitude + Elevation | 6 | 289.85 | 3.76 | 0.06 | -138.47 |
| EWPW-Wing.Chord-Breed-TR | | | | | |
| Temperature | 5 | 266.78 | 0 | 0.62 | -128.07 |
| Mean Daily Range + Temperature | 6 | 268.11 | 1.33 | 0.32 | -127.61 |
| Solar Radiation | 5 | 272.13 | 5.35 | 0.04 | -130.75 |
| Mean Daily Range + Solar Radiation | 6 | 274.18 | 7.39 | 0.02 | -130.64 |
| Mean Daily Range | 5 | 286.07 | 19.29 | 0 | -137.72 |
| 1 | 4 | 286.08 | 19.3 | 0 | -138.83 |
| EWPW-Wing.Chord-Winter-TR | | | | | |
| 1 | 4 | 286.08 | 0 | 0.42 | -138.83 |
| Mean Daily Range | 5 | 288.06 | 1.97 | 0.16 | -138.71 |
| Solar Radiation | 5 | 288.29 | 2.2 | 0.14 | -138.82 |
| Temperature | 5 | 288.29 | 2.2 | 0.14 | -138.83 |
| Mean Daily Range + Temperature | 6 | 290.31 | 4.23 | 0.05 | -138.7 |

| | | | | | |
|---|---|---|---|---|---|
| Mean Daily Range + Solar Radiation | 6 | 290.31 | 4.23 | 0.05 | -138.71 |
| Solar Radiation + Temperature | 6 | 290.55 | 4.46 | 0.05 | -138.82 |
| EWPW-Wing.Chord-Breed-Prod | | | | | |
| Precipitation | 5 | 264.67 | 0 | 0.73 | -127.02 |
| EVI + Precipitation | 6 | 266.71 | 2.04 | 0.27 | -126.9 |
| EVI | 5 | 283.78 | 19.11 | 0 | -136.57 |
| 1 | 4 | 286.08 | 21.41 | 0 | -138.83 |
| EWPW-Wing.Chord-Winter-Prod | | | | | |
| Precipitation | 5 | 283.65 | 0 | 0.56 | -136.51 |
| EVI + Precipitation | 6 | 285.74 | 2.08 | 0.2 | -136.42 |
| 1 | 4 | 286.08 | 2.43 | 0.17 | -138.83 |
| EVI | 5 | 287.88 | 4.22 | 0.07 | -138.62 |
| EWPW-Wing.Chord-Breed-Seas | | | | | |
| TemperatureCV | 5 | 272.73 | 0 | 0.75 | -131.05 |
| EVI CV + TemperatureCV | 6 | 274.96 | 2.22 | 0.25 | -131.03 |
| 1 | 4 | 286.08 | 13.35 | 0 | -138.83 |
| EVI CV | 5 | 286.8 | 14.07 | 0 | -138.08 |
| EWPW-Wing.Chord-Winter-Seas | | | | | |
| 1 | 4 | 286.08 | 0 | 0.49 | -138.83 |
| TemperatureCV | 5 | 287.56 | 1.48 | 0.23 | -138.46 |
| EVI CV | 5 | 288.02 | 1.93 | 0.19 | -138.69 |
| EVI CV + TemperatureCV | 6 | 289.42 | 3.33 | 0.09 | -138.26 |
| EWPW-Wing.Chord-NA-Mig.Dist | | | | | |
| Migratory Distance | 5 | 275.43 | 0 | 1 | -132.4 |
| 1 | 4 | 286.08 | 10.65 | 0 | -138.83 |
| EWPW-Mass-Breed-Geo | | | | | |

| | | | | | |
|---|---|---|---|---|---|
| Latitude + Longitude | 6 | 272.05 | 0 | 0.48 | -129.6 |
| Latitude + Elevation | 6 | 272.15 | 0.1 | 0.45 | -129.65 |
| Latitude | 5 | 275.88 | 3.83 | 0.07 | -132.64 |
| Longitude + Elevation | 6 | 287.36 | 15.31 | 0 | -137.26 |
| Longitude | 5 | 287.76 | 15.71 | 0 | -138.58 |
| Elevation | 5 | 289.5 | 17.45 | 0 | -139.45 |
| 1 | 4 | 302.89 | 30.84 | 0 | -147.25 |
| EWPW-Mass-Winter-Geo | | | | | |
| Latitude | 5 | 302.28 | 0 | 0.34 | -145.84 |
| 1 | 4 | 302.89 | 0.61 | 0.25 | -147.25 |
| Elevation | 5 | 303.95 | 1.67 | 0.15 | -146.67 |
| Latitude + Elevation | 6 | 304.36 | 2.08 | 0.12 | -145.75 |
| Longitude | 5 | 305.01 | 2.73 | 0.09 | -147.2 |
| Longitude + Elevation | 6 | 306.19 | 3.9 | 0.05 | -146.67 |
| EWPW-Mass-Breed-TR | | | | | |
| Temperature | 5 | 282.48 | 0 | 0.75 | -135.94 |
| Mean Daily Range + Temperature | 6 | 284.72 | 2.25 | 0.25 | -135.94 |
| Solar Radiation | 5 | 299.01 | 16.53 | 0 | -144.2 |
| Mean Daily Range + Solar Radiation | 6 | 301.25 | 18.78 | 0 | -144.2 |
| 1 | 4 | 302.89 | 20.41 | 0 | -147.25 |
| Mean Daily Range | 5 | 304.69 | 22.21 | 0 | -147.04 |
| EWPW-Mass-Winter-TR | | | | | |
| Solar Radiation | 5 | 301.76 | 0 | 0.35 | -145.58 |
| 1 | 4 | 302.89 | 1.13 | 0.2 | -147.25 |
| Solar Radiation + Temperature | 6 | 303.83 | 2.07 | 0.13 | -145.49 |
| Mean Daily Range + Solar Radiation | 6 | 303.93 | 2.17 | 0.12 | -145.54 |
| Temperature | 5 | 304.34 | 2.57 | 0.1 | -146.87 |
| Mean Daily Range | 5 | 304.94 | 3.17 | 0.07 | -147.17 |

| | | | | | |
|---|---|---|---|---|---|
| Mean Daily Range + Temperature | 6 | 306.47 | 4.7 | 0.03 | -146.81 |
| EWPW-Mass-Breed-Prod | | | | | |
| Precipitation | 5 | 282.94 | 0 | 0.69 | -136.17 |
| EVI + Precipitation | 6 | 284.5 | 1.56 | 0.31 | -135.83 |
| EVI | 5 | 301.38 | 18.44 | 0 | -145.39 |
| 1 | 4 | 302.89 | 19.96 | 0 | -147.25 |
| EWPW-Mass-Winter-Prod | | | | | |
| Precipitation | 5 | 299.83 | 0 | 0.47 | -144.62 |
| EVI + Precipitation | 6 | 300.52 | 0.69 | 0.34 | -143.84 |
| 1 | 4 | 302.89 | 3.06 | 0.1 | -147.25 |
| EVI | 5 | 303.22 | 3.39 | 0.09 | -146.31 |
| EWPW-Mass-Breed-Seas | | | | | |
| EVI CV + TemperatureCV | 6 | 281.13 | 0 | 0.82 | -134.14 |
| TemperatureCV | 5 | 284.26 | 3.13 | 0.17 | -136.83 |
| EVI CV | 5 | 291.34 | 10.21 | 0 | -140.37 |
| 1 | 4 | 302.89 | 21.76 | 0 | -147.25 |
| EWPW-Mass-Winter-Seas | | | | | |
| TemperatureCV | 5 | 301.68 | 0 | 0.47 | -145.54 |
| 1 | 4 | 302.89 | 1.21 | 0.26 | -147.25 |
| EVI CV + TemperatureCV | 6 | 303.76 | 2.08 | 0.17 | -145.46 |
| EVI CV | 5 | 304.7 | 3.02 | 0.1 | -147.05 |
| EWPW-Mass-NA-Mig.Dist | | | | | |
| Migratory Distance | 5 | 291.18 | 0 | 1 | -140.29 |
| 1 | 4 | 302.89 | 11.71 | 0 | -147.25 |
| EUNI-Wing.Chord-Breed-Geo | | | | | |
| 1 | 2 | 85.74 | 0 | 0.48 | -40.64 |
| Elevation | 3 | 88.09 | 2.34 | 0.15 | -40.56 |
| Longitude | 3 | 88.12 | 2.38 | 0.15 | -40.58 |
| Latitude | 3 | 88.13 | 2.39 | 0.15 | -40.59 |
| Longitude + Elevation | 4 | 90.75 | 5.01 | 0.04 | -40.54 |

| | | | | | |
|---|---|---|---|---|---|
| Latitude + Elevation | 4 | 90.76 | 5.02 | 0.04 | -40.55 |
| EUNI-Wing.Chord-Winter-Geo | | | | | |
| 1 | 2 | 85.74 | 0 | 0.37 | -40.64 |
| Elevation | 3 | 87.22 | 1.47 | 0.18 | -40.13 |
| Longitude | 3 | 87.55 | 1.81 | 0.15 | -40.29 |
| Latitude | 3 | 88.15 | 2.4 | 0.11 | -40.59 |
| Longitude + Elevation | 4 | 89.22 | 3.48 | 0.07 | -39.78 |
| Latitude + Elevation | 4 | 89.55 | 3.81 | 0.06 | -39.94 |
| Latitude + Longitude | 4 | 89.92 | 4.18 | 0.05 | -40.13 |
| Latitude + Longitude + Elevation | 5 | 91.33 | 5.59 | 0.02 | -39.36 |
| EUNI-Wing.Chord-Breed-TR | | | | | |
| 1 | 2 | 85.74 | 0 | 0.45 | -40.64 |
| Mean Daily Range | 3 | 88.07 | 2.33 | 0.14 | -40.56 |
| Solar Radiation | 3 | 88.13 | 2.39 | 0.14 | -40.59 |
| Temperature | 3 | 88.19 | 2.45 | 0.13 | -40.61 |
| Mean Daily Range + Temperature | 4 | 90.28 | 4.54 | 0.05 | -40.31 |
| Mean Daily Range + Solar Radiation | 4 | 90.64 | 4.9 | 0.04 | -40.49 |
| Solar Radiation + Temperature | 4 | 90.77 | 5.03 | 0.04 | -40.55 |
| Mean Daily Range + Solar Radiation + Temperature | 5 | 92.91 | 7.17 | 0.01 | -40.15 |
| EUNI-Wing.Chord-Winter-TR | | | | | |
| 1 | 2 | 85.74 | 0 | 0.44 | -40.64 |
| Temperature | 3 | 87.53 | 1.79 | 0.18 | -40.29 |
| Solar Radiation | 3 | 87.9 | 2.16 | 0.15 | -40.47 |
| Mean Daily Range | 3 | 88.22 | 2.48 | 0.13 | -40.63 |

| | | | | | |
|---|---|---|---|---|---|
| Solar Radiation + Temperature | 4 | 90.11 | 4.37 | 0.05 | -40.22 |
| Mean Daily Range + Temperature | 4 | 90.22 | 4.48 | 0.05 | -40.28 |
| EUNI-Wing.Chord-Breed-Prod | | | | | |
| 1 | 2 | 85.74 | 0 | 0.6 | -40.64 |
| EVI | 3 | 88.13 | 2.38 | 0.18 | -40.58 |
| Precipitation | 3 | 88.23 | 2.48 | 0.17 | -40.63 |
| EVI + Precipitation | 4 | 90.76 | 5.02 | 0.05 | -40.55 |
| EUNI-Wing.Chord-Winter-Prod | | | | | |
| 1 | 2 | 85.74 | 0 | 0.51 | -40.64 |
| EVI | 3 | 87.13 | 1.39 | 0.26 | -40.09 |
| Precipitation | 3 | 88.11 | 2.37 | 0.16 | -40.58 |
| EVI + Precipitation | 4 | 89.64 | 3.9 | 0.07 | -39.99 |
| EUNI-Wing.Chord-Breed-Seas | | | | | |
| 1 | 2 | 85.74 | 0 | 0.57 | -40.64 |
| EVI CV | 3 | 87.83 | 2.09 | 0.2 | -40.44 |
| TemperatureCV | 3 | 88.12 | 2.38 | 0.17 | -40.58 |
| EVI CV + TemperatureCV | 4 | 90.48 | 4.73 | 0.05 | -40.4 |
| EUNI-Wing.Chord-Winter-Seas | | | | | |
| EVI CV | 3 | 83.95 | 0 | 0.55 | -38.49 |
| 1 | 2 | 85.74 | 1.8 | 0.22 | -40.64 |
| EVI CV + TemperatureCV | 4 | 86.44 | 2.5 | 0.16 | -38.39 |
| TemperatureCV | 3 | 88.13 | 4.18 | 0.07 | -40.59 |
| EUNI-Wing.Chord-NA-Mig.Dist | | | | | |
| 1 | 2 | 85.74 | 0 | 0.76 | -40.64 |
| Migratory Distance | 3 | 88.05 | 2.31 | 0.24 | -40.55 |
| EUNI-Mass-Breed-Geo | | | | | |
| 1 | 2 | 97.03 | 0 | 0.39 | -46.32 |

| | | | | | |
|---|---|---|---|---|---|
| Longitude | 3 | 98.32 | 1.28 | 0.2 | -45.75 |
| Latitude | 3 | 98.59 | 1.55 | 0.18 | -45.88 |
| Elevation | 3 | 99.44 | 2.41 | 0.12 | -46.31 |
| Longitude + Elevation | 4 | 100.64 | 3.61 | 0.06 | -45.61 |
| Latitude + Elevation | 4 | 100.93 | 3.9 | 0.05 | -45.75 |
| EUNI-Mass-Winter-Geo | | | | | |
| Longitude | 3 | 96.1 | 0 | 0.32 | -44.63 |
| 1 | 2 | 97.03 | 0.94 | 0.2 | -46.32 |
| Longitude + Elevation | 4 | 98.02 | 1.92 | 0.12 | -44.3 |
| Latitude | 3 | 98.49 | 2.4 | 0.1 | -45.83 |
| Elevation | 3 | 98.52 | 2.42 | 0.1 | -45.85 |
| Latitude + Longitude | 4 | 98.55 | 2.45 | 0.09 | -44.56 |
| Latitude + Elevation | 4 | 100.47 | 4.38 | 0.04 | -45.52 |
| Latitude + Longitude + Elevation | 5 | 100.75 | 4.66 | 0.03 | -44.27 |
| EUNI-Mass-Breed-TR | | | | | |
| Solar Radiation | 3 | 95.32 | 0 | 0.35 | -44.25 |
| Mean Daily Range + Solar Radiation | 4 | 96.83 | 1.5 | 0.17 | -43.7 |
| 1 | 2 | 97.03 | 1.71 | 0.15 | -46.32 |
| Solar Radiation + Temperature | 4 | 97.39 | 2.06 | 0.13 | -43.98 |
| Temperature | 3 | 98.44 | 3.11 | 0.07 | -45.81 |
| Mean Daily Range | 3 | 98.51 | 3.19 | 0.07 | -45.84 |
| Mean Daily Range + Solar Radiation + Temperature | 5 | 99.62 | 4.3 | 0.04 | -43.7 |
| Mean Daily Range + Temperature | 4 | 100.89 | 5.56 | 0.02 | -45.73 |
| EUNI-Mass-Winter-TR | | | | | |
| Mean Daily Range | 3 | 95.68 | 0 | 0.46 | -44.43 |
| 1 | 2 | 97.03 | 1.36 | 0.24 | -46.32 |

| | | | | | |
|---|---|---|---|---|---|
| Mean Daily Range + Temperature | 4 | 98.26 | 2.58 | 0.13 | -44.42 |
| Solar Radiation | 3 | 99.21 | 3.54 | 0.08 | -46.19 |
| Temperature | 3 | 99.46 | 3.78 | 0.07 | -46.32 |
| Solar Radiation + Temperature | 4 | 101.76 | 6.09 | 0.02 | -46.17 |
| EUNI-Mass-Breed-Prod | | | | | |
| EVI | 3 | 96.91 | 0 | 0.32 | -45.04 |
| 1 | 2 | 97.03 | 0.12 | 0.3 | -46.32 |
| Precipitation | 3 | 97.39 | 0.48 | 0.25 | -45.28 |
| EVI + Precipitation | 4 | 98.63 | 1.72 | 0.13 | -44.6 |
| EUNI-Mass-Winter-Prod | | | | | |
| 1 | 2 | 97.03 | 0 | 0.57 | -46.32 |
| EVI | 3 | 99.17 | 2.14 | 0.2 | -46.17 |
| Precipitation | 3 | 99.46 | 2.43 | 0.17 | -46.32 |
| EVI + Precipitation | 4 | 101.53 | 4.49 | 0.06 | -46.05 |
| EUNI-Mass-Breed-Seas | | | | | |
| EVI CV | 3 | 96.27 | 0 | 0.42 | -44.72 |
| 1 | 2 | 97.03 | 0.76 | 0.29 | -46.32 |
| EVI CV + TemperatureCV | 4 | 98.18 | 1.91 | 0.16 | -44.37 |
| TemperatureCV | 3 | 98.69 | 2.41 | 0.13 | -45.93 |
| EUNI-Mass-Winter-Seas | | | | | |
| 1 | 2 | 97.03 | 0 | 0.49 | -46.32 |
| EVI CV | 3 | 98.37 | 1.34 | 0.25 | -45.77 |
| TemperatureCV | 3 | 99.35 | 2.32 | 0.16 | -46.26 |
| EVI CV + TemperatureCV | 4 | 100.25 | 3.22 | 0.1 | -45.41 |
| EUNI-Mass-NA-Mig.Dist | | | | | |
| 1 | 2 | 97.03 | 0 | 0.7 | -46.32 |
| Migratory Distance | 3 | 98.72 | 1.68 | 0.3 | -45.95 |

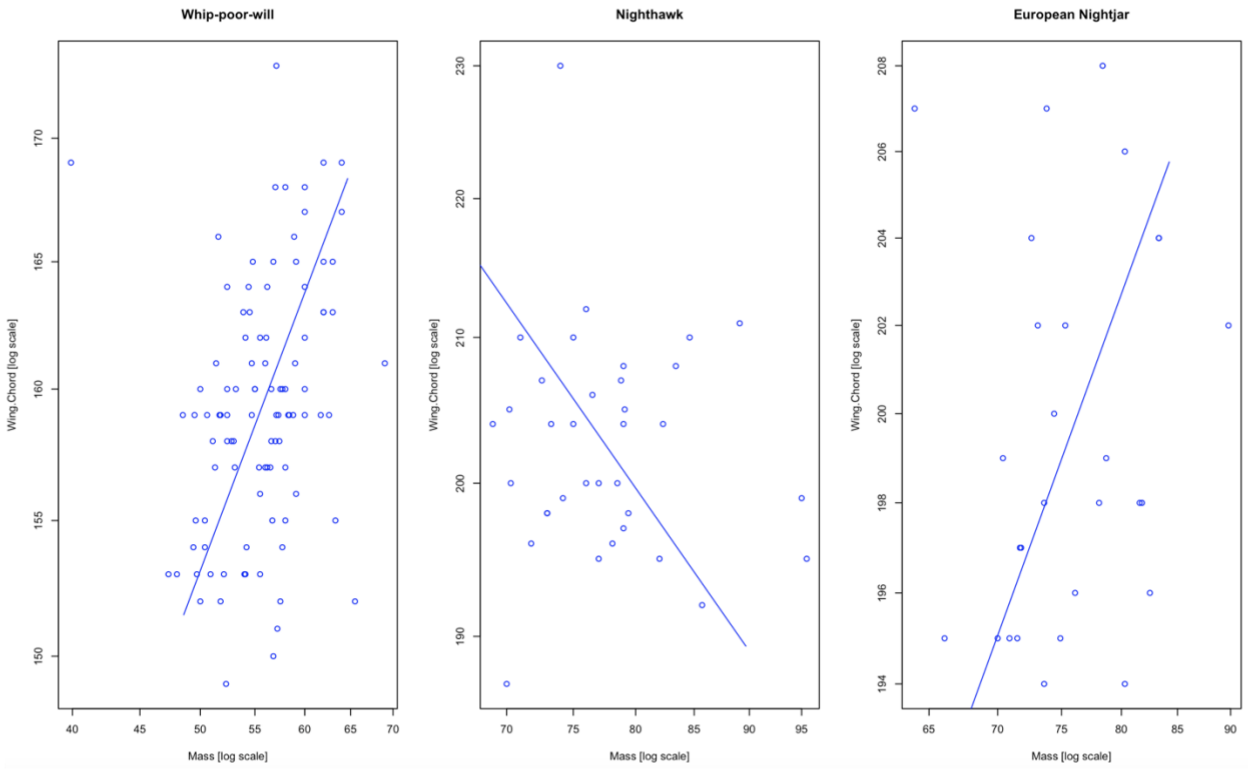


**Figure S1.** Allometric scaling theory provides the theoretical framework to link wing length and mass, where isometric scaling predicts that wing length (linear) increases proportionally with body mass (volume) to the one-third power. Whip-poor-wills and European nightjars exhibit near isometric scaling (scaling coefficient = 0.37 and 0.29, respectively). Scaling in nighthawks, on the other hand, differs substantially from isometric (scaling coefficient = -0.46). Scaling coefficients were determined and plots were produced using standardized major axis regression in the *smatr* package (Warton *et al.*, 2012).

### Common nighthawk breeding predictors

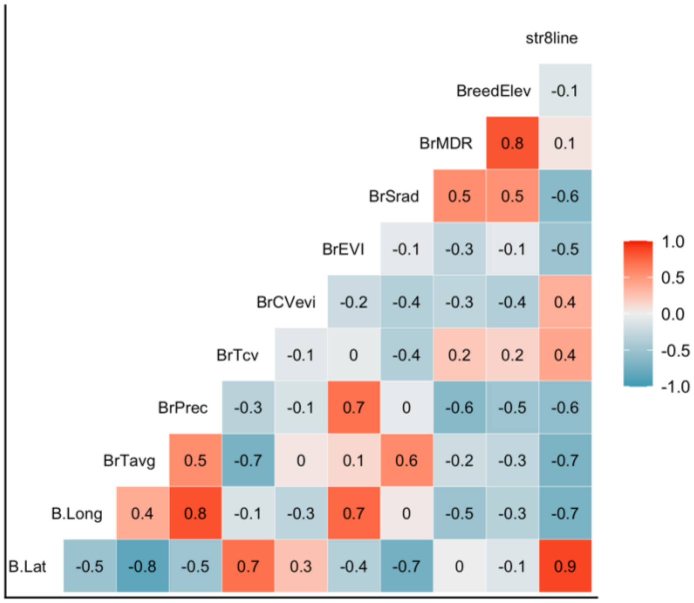


### Common nighthawk winter predictors

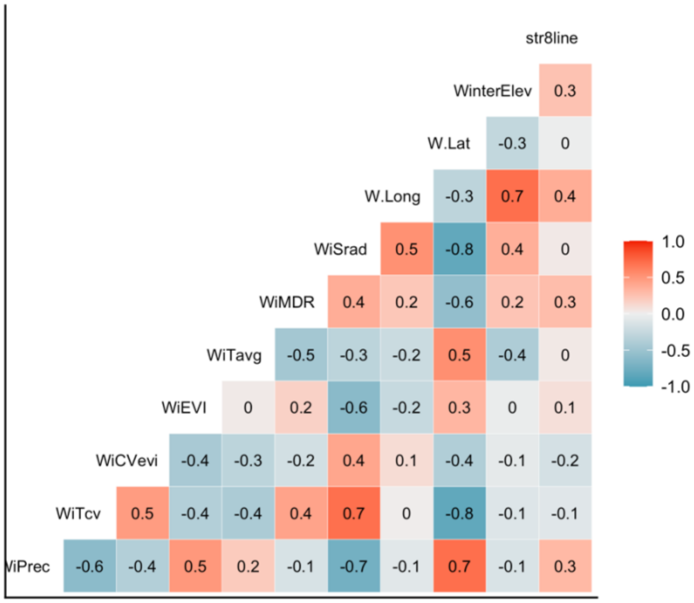

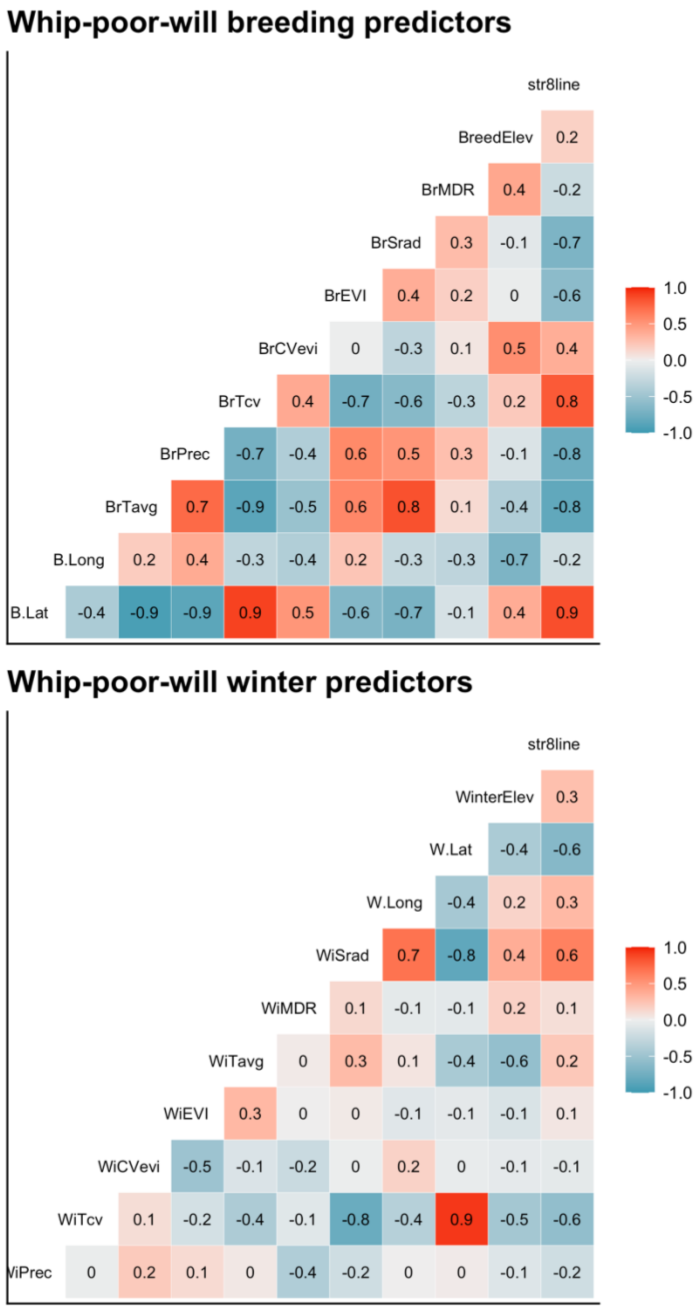


**Figure S2.** Correlation matrix plots of predictor variables for whip-poor-will and nighthawk on both the breeding and wintering grounds. We used the mass data set for both species as sample sizes were larger.
